# Isoliensinine Ameliorates Pathological Cardiac Hypertrophy After Aerobic Exercise by Activating Cytochrome c Oxidase Subunit 6A2

**DOI:** 10.1101/2025.07.07.663610

**Authors:** Shaolin Gong, Xia Feng, Xiaoting Jiang, Yaguang Bi, Fan Xia, Yongnan Fu, Xiaoping Peng, Zhiping Xiong, Xiang Wang

**Affiliations:** Department of Cardiology, The First Affiliated Hospital, Jiangxi Medical College, Nanchang University, Nanchang, Jiangxi, China; Academician Workstation of Cardiovascular Innovative Materials, Nanchang, Jiangxi, China; Jiangxi Hypertension Research Institute, Nanchang, Jiangxi, China; Jiangxi Key Laboratory of Neurological Diseases, Department of Cardiology, The First Affiliated Hospital, Jiangxi Medical College, Nanchang University, Nanchang, Jiangxi, China; Department of Cardiology, Zhongshan Hospital, Fudan University, Shanghai Institute of Cardiovascular Diseases

**Keywords:** Pathological myocardial hypertrophy, Aerobic exercise, Mitochondria, Isoliensinine, COX6A2

## Abstract

Pathological cardiac hypertrophy is an abnormal remodeling process of the heart under chronic pressure overload, characterized by myocardial fibrosis, apoptosis, and progressive decline in cardiac function, ultimately leading to heart failure or death. Although aerobic exercise, particularly swimming, has been shown to alleviate pathological cardiac hypertrophy, its molecular mechanisms remain incompletely understood. In this study, metabolomic and transcriptomic analyses revealed the molecular mechanisms underlying swimming-induced improvement of pathological cardiac hypertrophy. Results showed that swimming training significantly altered the level of isoliensinine in animals. Further studies demonstrated that isoliensinine exerts cardioprotective effects by regulating the expression of Cytochrome C Oxidase Subunit 6A2 (COX6A2), establishing for the first time the critical role of the Isoliensinine-COX6A2 signaling axis in cardiac function regulation. To validate this axis, a pathological cardiac hypertrophy model was constructed in mice through transverse aortic constriction (TAC). Both swimming training and isoliensinine intervention significantly improved cardiac structure and function. In vitro experiments using isoproterenol (ISO)-induced hypertrophy in H9C2 cells further confirmed the essential role of this signaling axis in suppressing pathological hypertrophy and maintaining mitochondrial homeostasis. Our findings demonstrate that swimming mitigates pathological cardiac hypertrophy by enhancing COX6A2 expression by isoliensinine, thereby improving mitochondrial function. This study not only expands the understanding of exercise-mediated mechanisms in pathological cardiac remodeling but also highlights the cardioprotective potential of isoliensinine and COX6A2 as therapeutic targets for preventing and treating heart failure.

## 1. Introduction

Pathological cardiac hypertrophy is a maladaptive response of the heart to sustained pressure overload, typically characterized by cardiomyocyte enlargement, extracellular matrix remodeling, and fibrosis ^1^. As pathological hypertrophy progresses, the heart gradually loses compensatory capacity, leading to systolic or diastolic dysfunction. This detrimental process is closely linked to increased risks of sudden cardiac death, heart failure, and cardiovascular mortality ^2^. Although current therapies, including pharmacological and device-based interventions, aim to reduce cardiac pressure overload, fewer than 50% of patients respond adequately due to various limitations, and effective molecularly targeted treatments remain scarce ^3^. Thus, exploring the regulatory mechanisms underlying pathological cardiac hypertrophy is critical for controlling the progression of cardiovascular diseases.

Since the mid-20th century, when Morris and Crawford first revealed the preventive effects of physical activity on cardiovascular diseases, aerobic exercise training is commonly accepted as a non-pharmacological intervention with significant benefits for cardiovascular health ^4,5^. Recent studies indicate that aerobic exercise effectively reduces cardiovascular issues like hypertension ^6^, insulin resistance ^7^, and provides cardioprotective effects ^4^. Specifically in heart failure management, cardiac rehabilitation programs based on exercise have been demonstrated to be effective in significantly reducing rehospitalization rates and all-cause mortality ^8^. Among aerobic exercise modalities, swimming training is particularly beneficial for cardiac health due to its low-impact nature and ability to modulate whole-body conditioning ^9^. Previous studies suggest that swimming not only enhances cardiovascular function but also reduces cardiac pressure overload and may partially reverse pathological cardiac hypertrophy ^10^. However, the exact molecular mechanisms behind how swimming training improves pathological cardiac hypertrophy are still not fully understood. Current evidence indicates that swimming mitigates pathological cardiac remodeling and regulates cardiac structure/function by promoting metabolic balance and enhancing antioxidant defenses ^11^.

The central role of metabolic imbalance and oxidative stress in the development of cardiac hypertrophy has been increasingly recognized ^12^. Under physiological conditions, cardiomyocytes rely heavily on fatty acid β-oxidation and glucose metabolism to maintain energy homeostasis. In pathological states, however, cardiac metabolism shifts toward glycolysis, accompanied by mitochondrial dysfunction (reduced ATP production), lipid buildup, and overproduction of reactive oxygen species (ROS) ^13^. This metabolic reprogramming not only directly impairs contractile function but also promotes pathological hypertrophy by activating signaling pathways like AMPK/mTOR ^14^ and HIF-1α ^15^. Concurrently, oxidative stress, a key downstream consequence of metabolic imbalance, exacerbates endoplasmic reticulum stress, inflammation, and apoptosis through damage to DNA, proteins, and lipids ^16,17^. Previous studies suggest that swimming training improves metabolic regulation and suppresses oxidative stress through pathways like AMPK/PGC-1α and Nrf2 ^18,19^. Nevertheless, the dynamic regulatory network by which swimming modulates metabolic balance and oxidative stress in pathological hypertrophy remains incompletely clear. Thus, elucidating the interplay between aerobic exercise and metabolic remodeling in pathological cardiac hypertrophy may offer novel strategies to overcome current therapeutic limitations.

Based on the aforementioned research, this study focuses on investigating the potential mechanisms by which swimming training suppresses pathological cardiac hypertrophy through metabolic remodeling. By establishing in vivo and in vitro pathological hypertrophy models and integrating metabolomics and transcriptomics, we aim to identify key molecular targets through which swimming regulates pathological cardiac hypertrophy, providing novel insights and strategies to reverse this detrimental process.

## 2. Materials and Methods

### 2.1 Establishment and Treatment of Pathological Cardiac Hypertrophy Mouse Model

Wild-type C57BL/6J mice were obtained from Cyagen Biosciences in Guangzhou, China. Mice were maintained in a standard SPF environment (temperature and humidity controlled, 12-hour light-dark cycle) with ad libitum access to water and food. All animal protocols were approved by the Animal Ethics Committee of the First Affiliated Hospital of Nanchang University (Approval No.: CDYFY-IACUC-202407QR148 and CDYFY-IACUC-202206QR008) and conducted in accordance with NIH guidelines. After a week of acclimation, the mice were randomly allocated to different experimental groups.

1) To investigate the effects of early exercise intervention on pathological cardiac hypertrophy, mice underwent a swimming training protocol as follows: A sterilized transparent glass tank (45 × 35 × 20 cm³) was filled with distilled water to a depth of 15 cm, maintained at 30 ± 1°C. Mice in the Swim group performed two 10-minute adaptation sessions on the first day (4–6 hours apart). The daily swimming duration was progressively increased by 10 minutes, reaching 90 minutes per session (twice daily) by Day 9, and sustained until Day 21. Control group (CON) mice were placed in an identical empty tank to simulate the procedure environment. After each session, Swim group mice were transferred using a mesh net, gently dried with absorbent gauze, and thoroughly dried with 37°C warm air before returning to standard housing alongside the CON group. Seven days after the final training session, TAC was performed to establish the pathological cardiac hypertrophy model.

2) TAC Surgery: Mice were administered isoflurane anesthesia, intubated, and connected to a ventilator. A thoracotomy was carried out through the second intercostal space to reveal the aortic arch. A 6-0 suture was tied around the aortic arch, with a 27G needle used as a spacer, followed by needle removal to create a stenotic lumen. The incision was closed with layered sutures. Four weeks post-TAC, mice were euthanized to collect blood and cardiac tissues. Blood samples were centrifuged to isolate serum, and ANP and BNP levels were quantified by enzyme-linked immunosorbent assay (ELISA). Cardiac tissues were processed for histopathological staining and protein analysis.

3) To validate the functional role of isoliensinine (ILS), exogenous ILS (HY-N0770, MCE, USA) was administered to assess its effects on cardiac structure and function in TAC mice. The experimental groups were as follows: Control: Sham surgery (thoracotomy without aortic constriction); ILS: Sham surgery + daily ILS (20 mg/kg/day, oral gavage); TAC: TAC surgery alone; TAC+ILS: TAC surgery + daily ILS (20 mg/kg/day, oral gavage).

### 2.2 Echocardiography

Cardiac performance was evaluated with a Vevo 3100 high-frequency ultrasound system (Canada). After depilation of the chest and induction of anesthesia (1.5–2% isoflurane), body temperature was maintained at ∼37 °C to reduce motion artifacts. A thin layer of conductive gel was applied to the probe, and parasternal short-axis M-mode images were captured at the level of the left-ventricular papillary muscles. From these recordings, we determined ejection fraction (EF), fractional shortening (FS), interventricular septal thickness in diastole (IVSd), left-ventricular posterior wall thickness in diastole (LVPWd), end-systolic and end-diastolic diameters (LVESd, LVEDd), as well as their respective volumes (LVESV, LVEDV). All measurements were analyzed using Vevo LAB software.

### 2.3 Histopathological Staining

Tissues were first fixed in 4% paraformaldehyde for 24 h, then subjected to graded dehydration, cleared in xylene, and embedded in paraffin. Embedded blocks were cut into 4–5 µm sections with a rotary microtome, after which slides were deparaffinized in xylene, rehydrated through a descending ethanol series, and rinsed in distilled water. Histological staining was carried out following manufacturers’ protocols: hematoxylin & eosin (G1120, Solarbio, China), Masson’s trichrome (G1340, Solarbio, China), and wheat germ agglutinin (WGA) fluorescence (GTX01502, GeneTex, USA).

### 2.4 Heart Weight/Body Weight Ratio and Heart Weight/Tibia Length Ratio

Following euthanasia, each animal’s terminal body weight was noted. Hearts were promptly removed, rinsed in ice-cold saline, blotted dry, and weighed to determine the wet heart weight (HW). The left tibia was then isolated, stripped of surrounding soft tissue, and its length—from the proximal joint surface to the distal ankle—measured with a digital caliper. Ratios were calculated as: HW/BW (mg/g) = wet heart weight (mg) ÷ final body weight (g); HW/TL (mg/mm) = wet heart weight (mg) ÷ tibia length (mm)

### 2.5 qPCR Analysis

Cardiac tissues (≈30 mg) were homogenized in lysis buffer, while H9C2 cell pellets were lysed directly with the same buffer. Total RNA was isolated using a mei5bio kit (Beijing, China), and its yield and purity were assessed via NanoDrop (Thermo Fisher Scientific, USA). A 500 ng aliquot of RNA was reverse-transcribed into cDNA with a Vazyme kit (China). qPCR was then carried out using gene-specific primers (Genewiz, Suzhou, China; see Table 1) in a conventional reaction setup.

**Table 1.**
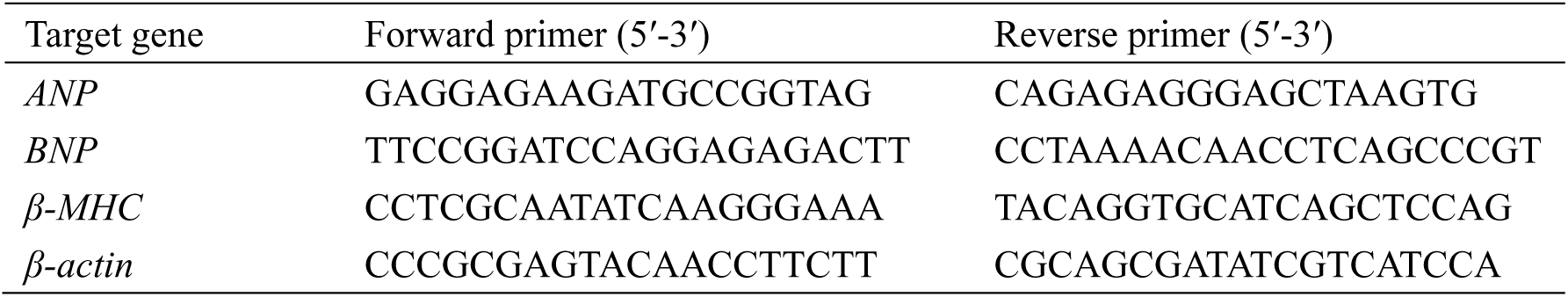
Sequence information of the primers used for quantitative reverse transcription-PCR.

### 2.6 TUNEL Assay

Cardiac samples were fixed in 4% paraformaldehyde for 24–48 h, then dehydrated through an ethanol series and embedded in paraffin. Four-micron sections were cut, dewaxed, and rehydrated, after which antigens were unmasked in citrate buffer. Slides were permeabilized with 20 µg/mL Proteinase K at 37 °C for 15 min, rinsed three times in PBS, and then treated with TUNEL reaction mix (Beyotime, China) at 37 °C in the dark for 30 min. After a further three PBS washes, nuclei were stained with DAPI for 5 min. Finally, coverslips were mounted with anti-fade medium, and images were acquired by fluorescence microscopy for analysis.

### 2.7 Measurement of Cardiomyocyte Contractile Function

Freshly isolated mouse ventricular myocytes were placed in a 37 °C perfusion chamber filled with Tyrode’s solution (1.8 mM Ca²⁺) and allowed to stabilize for 10 minutes. Mechanical function was recorded in real time with the IonOptix Myocam system (IonOptix Inc., Milton, MA, USA) under electrical field stimulation (0.5 Hz, 5 ms pulse). The following metrics were measured: resting cell length; +dL/dt (rate of contraction); peak shortening amplitude; −dL/dt (rate of relaxation); time to peak shortening (TPS); and time to 90 % relengthening (TR₉₀).

### 2.8 ELISA Assay

Serum ANP and BNP levels were measured through ELISA. In accordance with the manufacturer’s guidelines (Elabscience Biotechnology, Wuhan, China), samples underwent coating, blocking, and incubation. A microplate reader was used to measure absorbance at 450 nm (OD=450 nm), enabling the calculation of target substance concentrations in the samples.

### 2.9 Metabolomic and Transcriptomic Analyses

Metabolomic and transcriptomic analyses were conducted by Oebiotech Co., Ltd. (Shanghai, China) and Biotree Co., Ltd. (Shanghai, China), respectively. For metabolomics analysis, samples were extracted with 80% methanol, vortexed and centrifuged, then assessed through ultra-high-performance liquid chromatography combined with quadrupole time-of-flight mass spectrometry (UHPLC-QTOF-MS). The mobile phase was gradient-eluted, with full scans in both positive/negative ion modes (m/z 50-1000). Raw data underwent peak extraction, metabolite annotation (based on HMDB/METLIN databases), and KEGG pathway enrichment analysis.

For RNA sequencing, total RNA was isolated from cardiac tissues using the TRIzol method. Strand-specific libraries were constructed and sequenced on the Illumina NovaSeq 6000 platform. Quality-filtered reads underwent alignment to the mouse genome using STAR. DESeq2 was employed to identify differentially expressed genes (FDR < 0.05), followed by analysis and visualization through KEGG and DAVID databases.

### 2.10 Determination of Mitochondrial Oxygen Consumption Rate (OCR)

Cardiomyocyte mitochondrial function was measured with the Seahorse XFp Mitochondrial Stress Test Kit (Agilent, USA). In brief, hearts from anesthetized mice were excised, and cardiomyocytes were isolated via calcium-free enzymatic digestion. Cells were plated on poly-lysine-coated Seahorse 96-well plates in XF assay medium and equilibrated at 37 °C without CO₂ for 1 hour. During the assay, the XF Analyzer injected oligomycin, FCCP, and rotenone/antimycin A in sequence to determine basal respiration, ATP-linked respiration, maximal respiration, and spare respiratory capacity. Data were processed in Wave software to quantify mitochondrial respiratory parameters.

### 2.11 Transmission Electron Microscope (TEM) Analysis

Cardiac specimens were immediately fixed in ice-cold glutaraldehyde, post-fixed with osmium tetroxide, then dehydrated through graded ethanol or acetone and embedded in epoxy resin. Ultrathin sections (50–70 nm) were cut, stained with uranyl acetate and lead citrate, and examined by transmission electron microscopy. Mitochondrial ultrastructure—such as cristae organization, matrix density, and membrane integrity—was then evaluated.

### 2.12 ROS&Mito SOX Detection Experiment

Fresh or frozen cardiac tissue sections (5-10 μm thickness) were incubated in the dark with DHE probe (Yeasen, China) or MitoSOX Green probe (Thermo Fisher Scientific, USA) at 37°C for 30-60 minutes. After washing to remove unbound probes, sections were briefly fixed with 4% paraformaldehyde, mounted with antifade mounting medium, and stored protected from light. Fluorescence intensity of ROS (red fluorescence, DHE) and MitoSOX (green fluorescence) was observed under a fluorescence microscope to evaluate superoxide anion levels in the cytoplasm and mitochondria.

### 2.13 Cell Treatment

To validate the anti-cardiac hypertrophic effects of ILS in vitro, H9C2 cells (Procell, Wuhan, China) were treated with isoprenaline (ISO, HY-B0468, MCE, USA) to establish a cellular hypertrophy model. Cells were maintained under standard culture conditions (high-glucose DMEM medium supplemented with 10% fetal bovine serum and 1% penicillin/streptomycin, 37°C, 5% CO₂) and divided into four groups: CON group (untreated control), ISO group (40 μM ISO, 24 h), ILS group (5 μM ILS, 24 h), ISO+ILS group (co-treatment with 40 μM ISO and 5 μM ILS, 24 h).

### 2.14 si-RNA transfection

H9C2 cells at 50% confluence were plated in 6-well plates and transfected with 50–100 nM COX6A2-specific siRNA or control siRNA (Genepharma, Shanghai, China) using a liposomal reagent. Six hours post-transfection, the medium was replaced with fresh complete medium, and cells were cultured for an additional 48 h. Knockdown efficiency was verified by qPCR and Western blot.

### 2.15 CCK8 Assay

H9C2 cells were plated in 96-well plates at ∼50% confluence and allowed to attach for 24 h. They were then assigned to a control group (standard medium) or to ILS treatment groups receiving different ILS concentrations for 24 h. After treatment, 10 µL of CCK-8 reagent (MCE, USA) was added to each well and incubated in the dark for 1 h. Absorbance at 450 nm was measured on a microplate reader, with cell-free wells used to correct background. Cell viability was calculated from the corrected OD values to evaluate ILS cytotoxicity.

### 2.16 Phalloidin staining

H9C2 cells were fixed in 4% paraformaldehyde, then permeabilized with 0.1% Triton X-100. Samples were incubated in the dark with FITC-phalloidin (Yeasen, China) at room temperature for 30 min, and excess dye was removed by PBS rinses. Nuclei were counterstained with DAPI, coverslips mounted using anti-fade medium, and F-actin architecture was visualized and photographed under a fluorescence microscope.

### 2.17 Western Blot Analysis

Samples were lysed in RIPA buffer (Solarbio, China) to extract total protein, and concentrations were measured by BCA assay. Equal amounts (30–50 µg) of protein were separated on 10% SDS–PAGE gels and transferred onto PVDF membranes. After blocking with 5% BSA for 1 h at room temperature, membranes were incubated overnight at 4 °C with primary antibodies against Caspase-3, Bax, Bcl-2, COX6A2, and β-actin (all 1:1 000, Proteintech, China). Following washes in TBST, membranes were probed with HRP-conjugated secondary antibodies (1:10 000, Proteintech, China) for 1 h at room temperature. Bands were visualized using ECL substrate, and the intensity of each target band was quantified relative to β-actin using ImageJ.

### 2.18 Statistical Analysis

Data are expressed as mean ± standard error of the mean (SEM). Statistical analysis was performed using an unpaired Student’s t-test for comparisons between two groups, while one-way ANOVA followed by Tukey’s post hoc test was used for comparisons among multiple groups. For experiments involving two independent variables, two-way ANOVA with Bonferroni correction was applied. Analyses were carried out using GraphPad Prism 10.0, and differences were considered statistically significant at *p* < 0.05.

## 3. Result

### 3.1 Swimming Training Attenuates TAC-Induced Pathological Cardiac Hypertrophy

To investigate the effects of aerobic exercise on pathological cardiac hypertrophy, a mouse swimming training model was employed. As shown in Figure 1A, after completing the swimming protocol, mice underwent transverse aortic constriction (TAC) or sham surgery. Echocardiographic assessment of cardiac function (Figure 1B-G) revealed that there was no significant difference in heart rate among the groups. However, compared to the Sham group, TAC-operated mice exhibited significant increases in LVIDd and LVIDs, along with markedly reduced EF, indicating TAC-induced left ventricular dilation and impaired systolic function. Further analysis demonstrated that the TAC group had significantly elevated heart weight-to-body weight ratio (HW/BW) and heart weight-to-tibia length ratio (HW/TL) compared to the Sham group, confirming pathological cardiac hypertrophy (Figure 1H-I). Molecular analysis showed that mRNA levels of hypertrophy markers (ANP and BNP) were significantly upregulated in the TAC group (Figure 1J-K).

**Figure 1.**
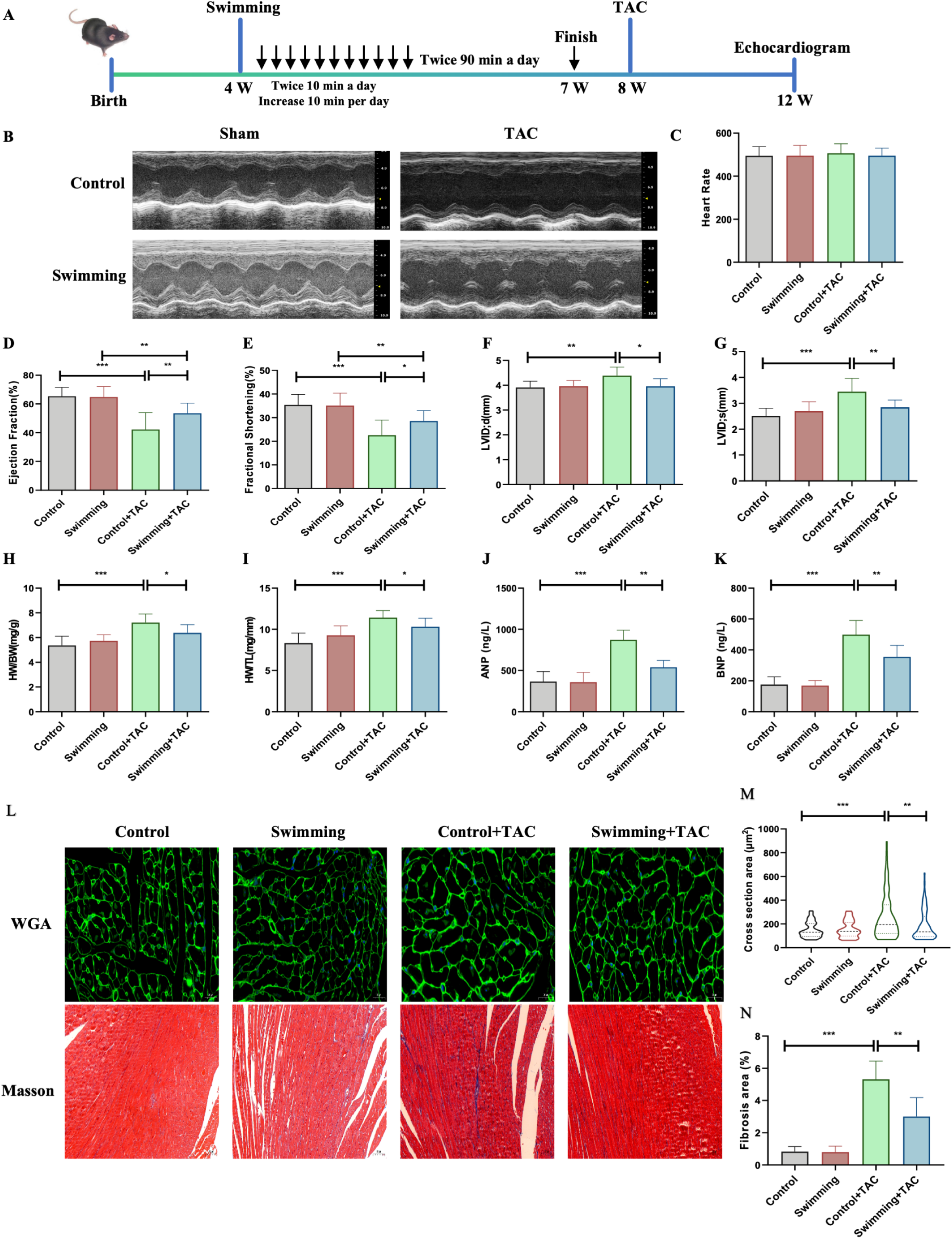
Swimming training mitigates TAC-induced pathological cardiac hypertrophy. (A) Experimental timeline: Mice underwent 3 weeks of swimming training followed by transverse aortic constriction (TAC) or sham surgery. Cardiac functional and molecular analyses were performed 4 weeks post-surgery. (B-G) Representative echocardiographic images and quantitative parameters of cardiac structure and function. TAC-operated mice exhibited ventricular dilation and impaired systolic function, which were ameliorated by swimming. (H-I) Heart weight-to-body weight ratio (HW/BW) and heart weight-to-tibia length ratio (HW/TL). (J-K) mRNA levels of hypertrophy markers ANP and BNP, normalized to β-actin. (L-N) Histopathological analysis. L: Representative images of wheat germ agglutinin (WGA) staining (green: cytoplasmic membrane; blue: nucleus) and Masson’s trichrome staining (blue: collagen; red: cardiomyocytes). M: Quantitative cardiomyocyte cross-sectional area. N: Percentage of collagen deposition area. Scale bars: 20 μm. Data presented as mean ± SEM (n = 6). *P < 0.05; **P < 0.01; ***P < 0.001; ****P < 0.0001.

Histopathological evaluation revealed that wheat germ agglutinin (WGA) staining displayed a significant increase in cardiomyocyte cross-sectional area in the TAC group, indicative of cellular hypertrophy (Figure 1L-M). Masson’s trichrome staining showed a higher percentage of collagen deposition in the myocardial interstitium of TAC mice, suggesting aggravated fibrosis (Figure 1N). Notably, the Swim+TAC group exhibited significant improvements in left ventricular remodeling, systolic dysfunction, and abnormal expression of hypertrophy-related genes/pathological markers compared to the TAC group (Figure 1N). In conclusion, TAC surgery successfully induced pathological cardiac hypertrophy with fibrosis in mice, while swimming training significantly ameliorated these pathological alterations.

### 3.2 Swimming Training Attenuates TAC-Induced Apoptosis and Contractile Dysfunction

To further validate the cardioprotective benefits of swimming, cardiomyocyte apoptosis and contractile function were evaluated. In the TUNEL assay (Figure 2A-B), the TAC group exhibited a significant increase in apoptosis compared to the Control group, whereas swimming (Swim+TAC group) markedly reduced apoptotic activity. Assessment of cardiomyocyte contractile dynamics (Figure 2C-H) revealed that TAC-operated mice showed significant increases in Resting Cell Length and TR90 (time to 90% recovery of resting length), while +dL/dt, -dL/dt, Peak Shortening, and TPS (time to peak shortening) were significantly decreased. Conversely, the Swim+TAC group demonstrated reversed trends in these parameters compared to the TAC group. These findings suggest that swimming training effectively improves TAC-impaired cardiomyocyte contractile function.

**Figure 2.**
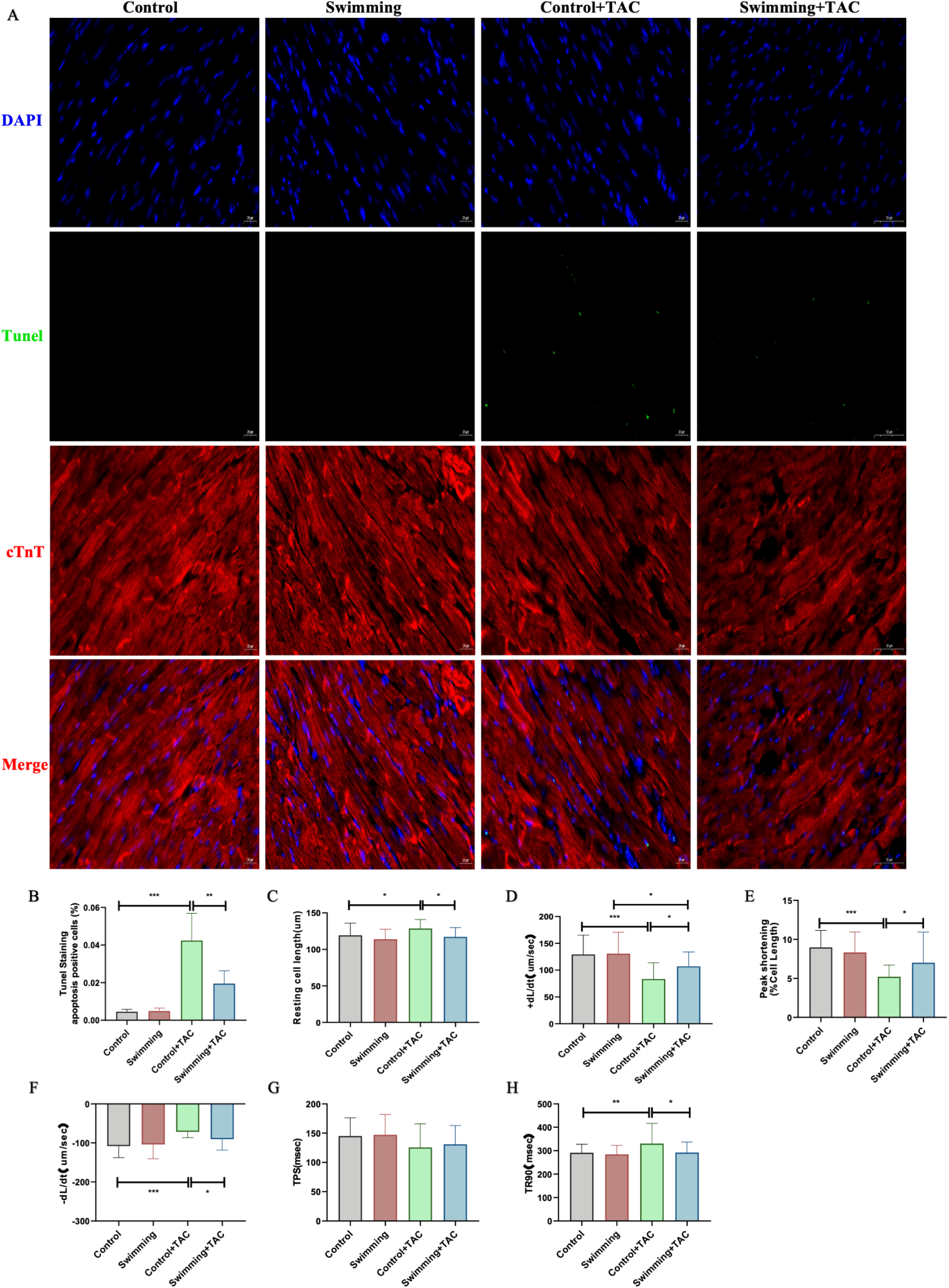
Swimming training attenuates TAC-induced cardiomyocyte apoptosis and contractile dysfunction. (A) Representative TUNEL staining images (green: apoptotic cells; blue: nuclei) in cardiac tissue sections. (B) Quantitative analysis of TUNEL-positive cells, expressed as percentage of total nuclei. Cardiomyocyte contractile dynamics assessed by IonOptix Myocam system: (C) Resting Cell Length (static cell length at baseline); (D) +dL/dt (maximal rate of contraction); (E) Peak Shortening (maximal shortening amplitude during contraction); (F) -dL/dt (maximal rate of relaxation); (G) TPS (time to peak shortening); (H) TR90 (time to 90% recovery of resting length). Data presented as mean ± SEM (n = 6). *\*P <* 0.05; *\*\*P <* 0.01; *\*\*\*P <* 0.001; *\*\*\*\*P <* 0.0001.

### 3.3 Isoliensinine Attenuates TAC-Induced Pathological Cardiac Hypertrophy

To investigate the mechanism underlying swimming-mediated cardioprotection, metabolomic assays was performed in Swim-trained and control mice (Figure 3A-B). Sequencing results highlighted a swimming-induced upregulation of isoliensinine, a bioactive compound. To validate its role, isoliensinine was administered exogenously to TAC-operated mice (Figure 3C). Echocardiography demonstrated that the Isoliensinine+TAC group exhibited significantly reduced LVIDd and improved EF compared to the TAC group, indicating alleviated ventricular dilation and systolic dysfunction (Figure 3D-I). Further analysis showed that the Isoliensinine+TAC group had significantly lower HW/BW and HW/TL than the TAC group (Figure 3J-K), confirming attenuated pathological hypertrophy. ELISA revealed markedly reduced serum levels of ANP and BNP in the Isoliensinine+TAC group (Figure 3L-M). Consistently, mRNA expression of hypertrophy markers ANP and BNP was significantly downregulated in the Isoliensinine+TAC group compared to the TAC group (Figure 3N).

**Figure 3.**
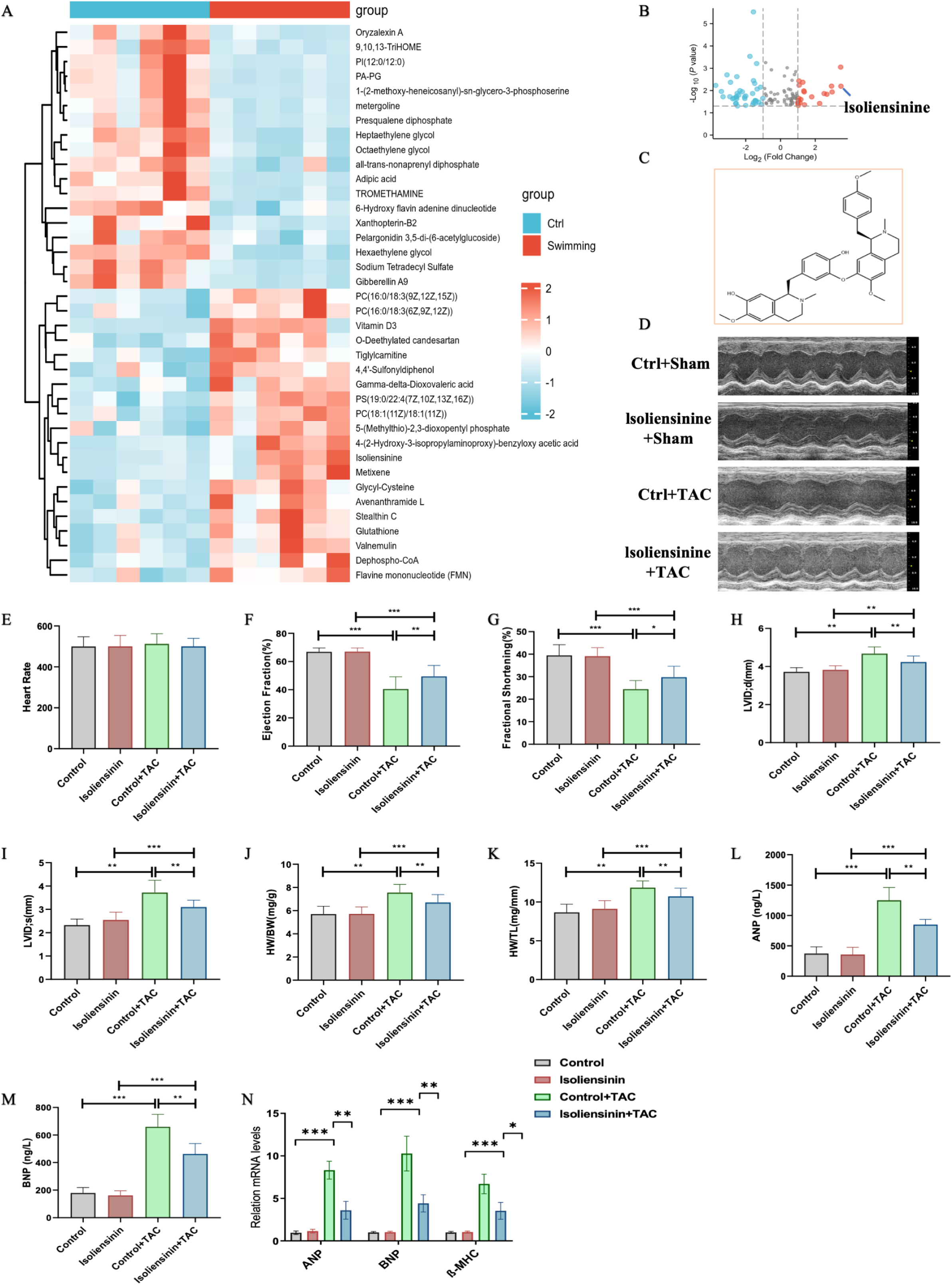
Isoliensinine mediates swimming-induced cardioprotection. Metabolomic profiling of cardiac tissues: (A) Heatmap of differentially expressed metabolites between Swim and control groups; (B) Volcano plot highlighting isoliensinine (red dot) as a key upregulated metabolite in the Swim group (log2 fold change >1.5, p < 0.05). (C) Chemical structure of isoliensinine, a bioactive alkaloid derived from Nelumbo nucifera. Echocardiographic assessment of cardiac function: (D) Representative images of left ventricular morphology; (E) Heart rate; (F) Ejection fraction (EF); (G) fractional shortening (FS); (H-I) Left ventricular end-diastolic diameter (LVIDd) and end-systolic diameter (LVIDs). Data presented as mean ± SEM (n = 10/group). P < 0.05 was considered to indicate a statistically significant difference. (J-K) Heart weight-to-body weight ratio (HW/BW) and heart weight-to-tibia length ratio (HW/TL). Isoliensinine significantly attenuated TAC-induced cardiac hypertrophy. (L-M) Serum levels of ANP and BNP measured by ELISA. (N) qPCR analysis of hypertrophy markers (ANP, BNP, β-MHC). mRNA levels were normalized to β-actin. Data presented as mean ± SEM (n = 6), *\*P <* 0.05; *\*\*P <* 0.01; *\*\*\*P <* 0.001; *\*\*\*\*P <* 0.0001.

### 3.4 Isoliensinine Attenuates TAC-Induced Myocardial Fibrosis and Apoptosis

Further validation of isoliensinine’s effects on pathological cardiac hypertrophy was conducted. Histopathological analysis (Figure 4A-C) demonstrated that WGA staining revealed a significantly smaller cardiomyocyte cross-sectional area in the Isoliensinine+TAC group compared to the TAC group, indicating alleviated cellular hypertrophy. Masson’s trichrome staining showed a lower percentage of collagen deposition in the myocardial interstitium of the Isoliensinine+TAC group, suggesting reduced fibrosis. Additionally, TUNEL staining confirmed significantly reduced apoptosis in the Isoliensinine+TAC group compared to the TAC group (Figure 4D-E). Collectively, these results demonstrate that isoliensinine effectively mitigates TAC-induced cardiomyocyte hypertrophy, myocardial fibrosis, and apoptosis.

**Figure 4.**
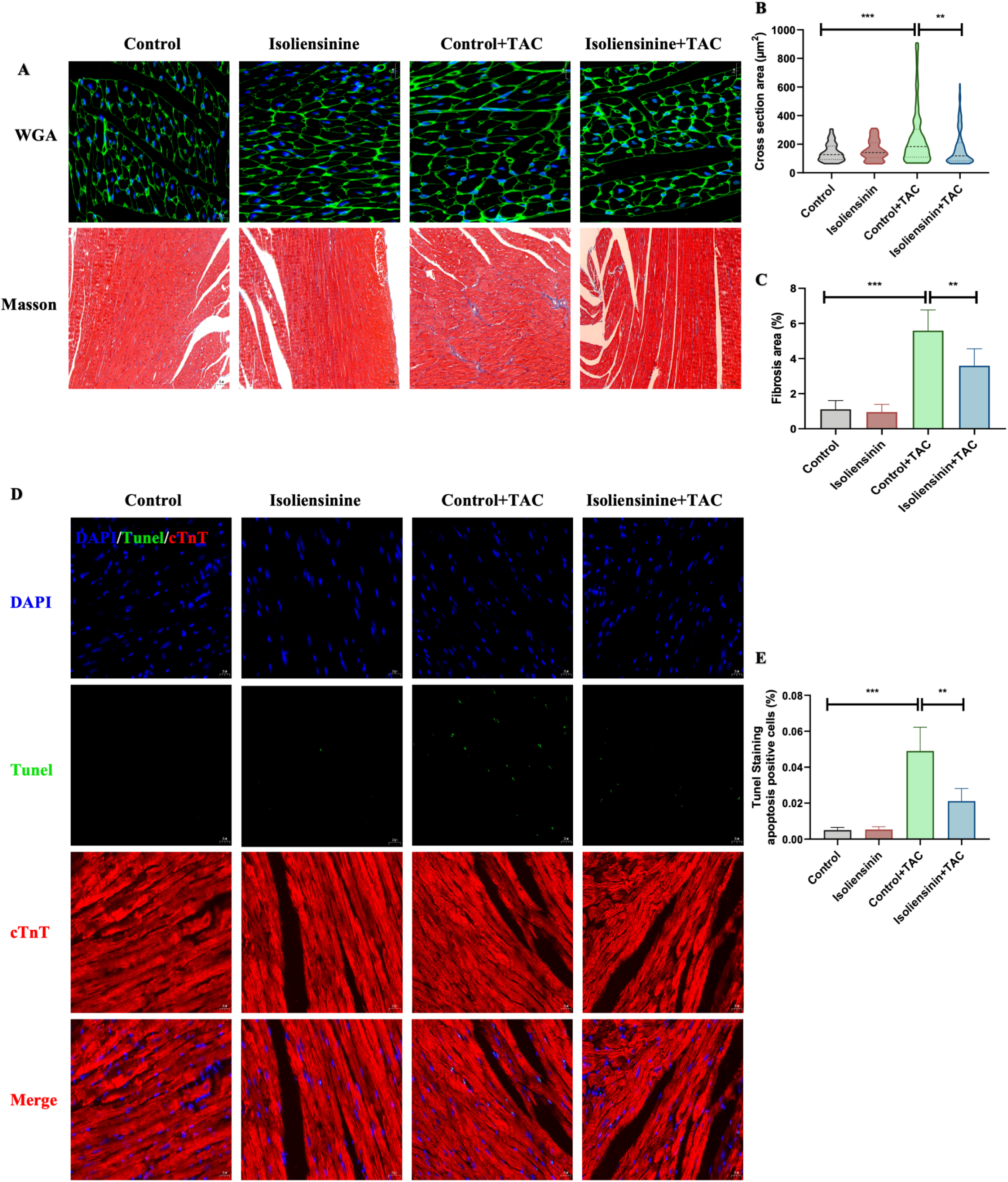
Isoliensinine alleviates TAC-induced cardiomyocyte hypertrophy, fibrosis, and apoptosis. (A) Representative images of Wheat Germ Agglutinin (WGA) staining (green: cell membranes; blue: nuclei) and Masson’s trichrome staining (blue: collagen; red: cardiomyocytes). (B) Quantitative analysis of cardiomyocyte cross-sectional area (CSA) from WGA staining. (C) Quantitative analysis of collagen deposition area (%) from Masson’s trichrome staining. (D) Representative TUNEL staining images (green: apoptotic cells; blue: nuclei). (E) Quantification of TUNEL-positive cells. Scale bars: 20 μm (A, D). Data presented as mean ± SEM (n = 6), *\*P <* 0.05; *\*\*P <* 0.01; *\*\*\*P <* 0.001; *\*\*\*\*P <* 0.0001.

### 3.5 Isoliensinine Improves Mitochondrial Respiratory Function

Oxygen consumption rate (OCR) was measured using the Seahorse XF Analyzer to explore the effects of swimming on mitochondrial respiration. Results showed that TAC surgery reduced OCR in cardiomyocytes, with consistent decreases in basal respiration, ATP production, maximal respiration, and spare respiratory capacity (Figure 5A-B). Isoliensinine increased OCR and reversed TAC-induced impairments in mitochondrial respiratory function, indicating enhanced mitochondrial respiration. Transmission electron microscopy (TEM) revealed structural abnormalities in mitochondria of TAC mice, including reduced cristae density and mitochondrial swelling/fusion (Figure 5C). These pathological changes were significantly ameliorated in the Isoliensinine+TAC group. Collectively, Isoliensinine improved both mitochondrial ultrastructure and respiratory function compromised by TAC.

**Figure 5.**
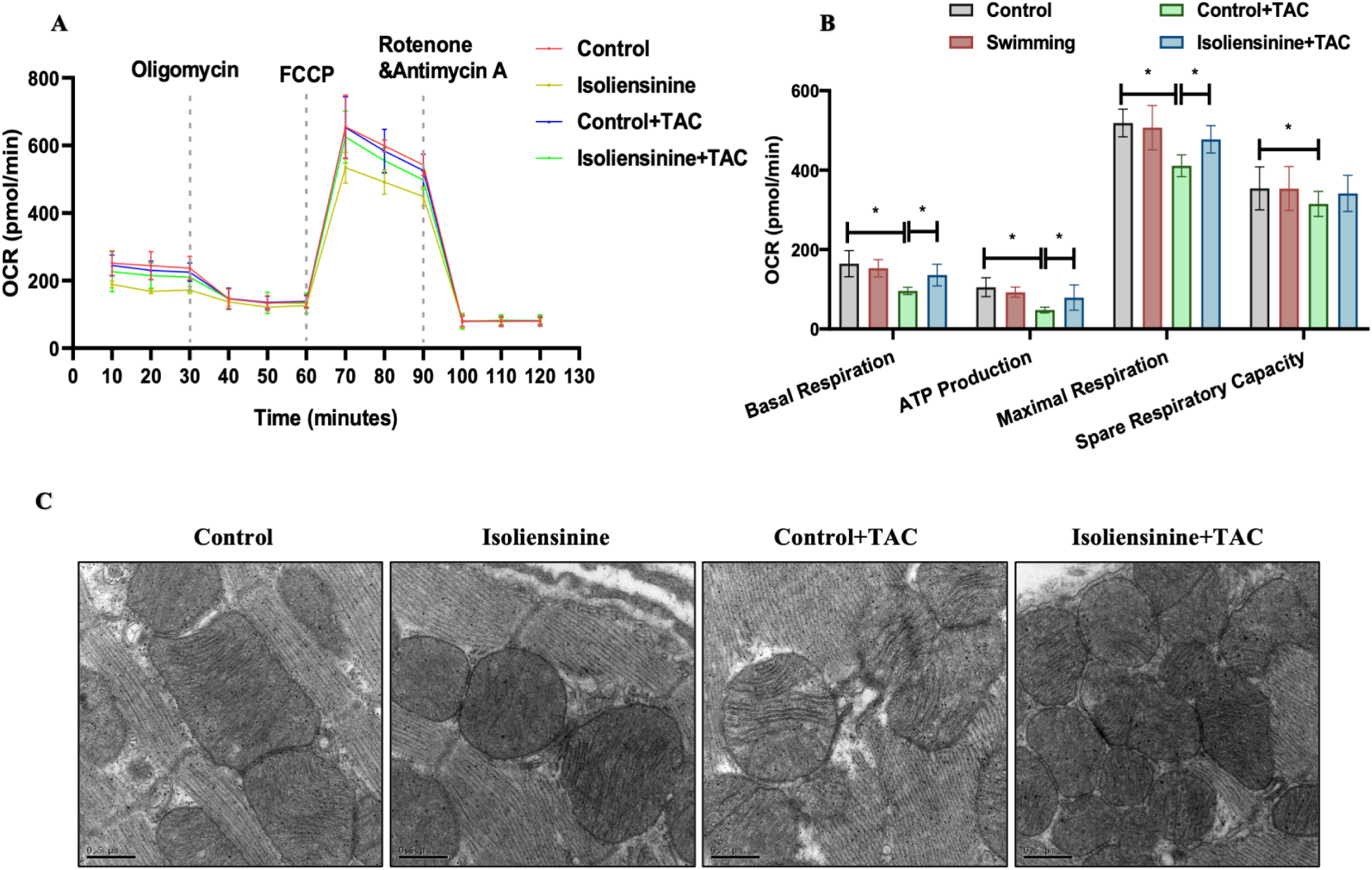
Isoliensinine restores mitochondrial respiratory function and ultrastructure in TAC-operated mice. (A) Mitochondrial oxygen consumption rate (OCR) measured using the Seahorse XFp Analyzer. Representative OCR trace illustrating mitochondrial respiratory parameters under sequential injections of oligomycin, FCCP, and rotenone/antimycin A. (B) Quantification of mitochondrial respiratory parameters: Basal Respiration (baseline OCR), ATP Production (OCR linked to ATP synthesis), Maximal Respiration (FCCP-uncoupled maximal OCR), and Spare Respiratory Capacity (maximal OCR minus basal OCR). (C) Transmission electron microscopy (TEM) images of cardiac mitochondria. Data presented as mean ± SEM (n = 6). *\*P <* 0.05; *\*\*P <* 0.01; *\*\*\*P <* 0.001; *\*\*\*\*P <* 0.0001.

### 3.6 Isoliensinine Attenuates ISO-Induced H9C2 Cell Injury

In addition, in vitro experiments were conducted using ISO-induced H9C2 cell hypertrophy to validate the effects of isoliensinine. A CCK-8 assay confirmed that 5 μM isoliensinine had no cytotoxic effects on cell viability (Figure 6A). qPCR analysis revealed that co-treatment with ISO and ILS (ISO+ILS group) significantly downregulated hypertrophic markers (ANP, BNP, and β-MHC) compared to the ISO group (Figure 6B-D). Phalloidin staining further demonstrated a marked reduction in cell surface area in the ISO+ILS group versus the ISO group, confirming ILS-mediated attenuation of ISO-induced cellular hypertrophy (Figure 6E-F). Fluorescence probes were used to assess oxidative stress levels. The ISO+ILS group exhibited significantly lower intracellular ROS and MitoSOX levels compared to the ISO group, indicating that ILS reduces ISO-triggered reactive oxygen species (ROS) generation and alleviates oxidative stress (Figure 6G-I). Western blot analysis of apoptosis-related proteins showed that the ISO group displayed upregulated Caspase3 and Bax expression and downregulated Bcl-2 expression compared to the control group. Conversely, the ISO+ILS group exhibited opposite trends relative to the ISO group, demonstrating ILS-mediated suppression of ISO-induced apoptosis (Figure 6J-M).

**Figure 6.**
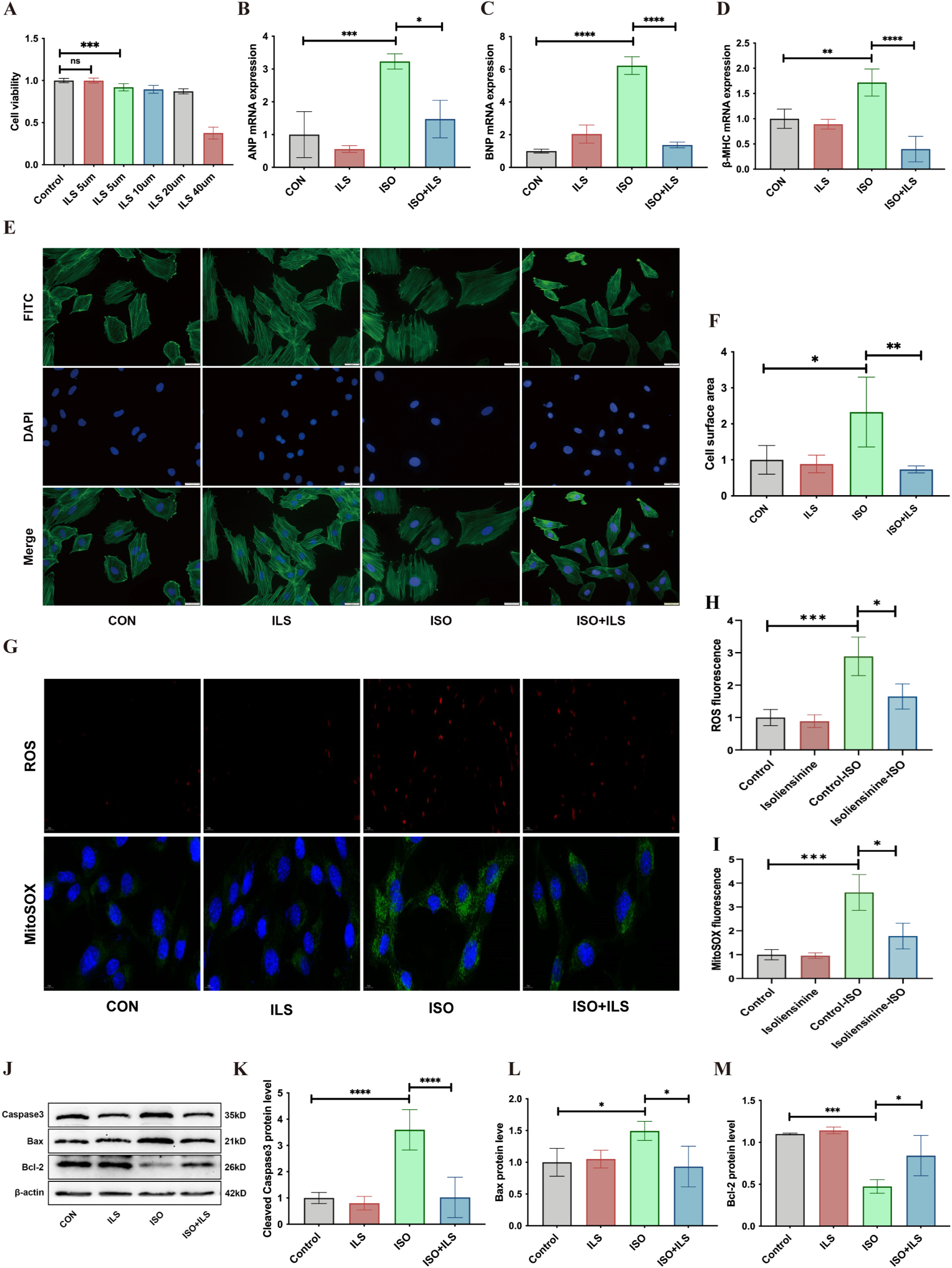
Isoliensinine attenuates ISO-induced H9C2 cell hypertrophy, oxidative stress, and apoptosis. (A) CCK-8 assay assessing cytotoxicity of isoliensinine (ILS). Cell viability remained unaffected at 5 μM ILS, confirming its non-toxic concentration for subsequent experiments. (B-D) qPCR analysis of hypertrophic markers: ANP, BNP, and β-MHC. mRNA levels were normalized to β-actin. (E-F) Phalloidin staining (red: F-actin; blue: nuclei) and quantitative analysis of cell surface area. (G-I) Fluorescence imaging and quantification of intracellular ROS (DHE probe, red fluorescence) and mitochondrial superoxide (MitoSOX Green probe, green fluorescence). (J-M) Western blot analysis of apoptosis-related proteins: Caspase3, Bax, Bcl-2, and β-actin (loading control). Data presented as mean ± SEM (n ≥ 3), *\*P <* 0.05; *\*\*P <* 0.01; *\*\*\*P <* 0.001; *\*\*\*\*P <* 0.0001.

### 3.7 Isoliensinine Exerts Cardiomyocyte Protection Through Targeting COX6A2

To further clarify the mechanism underlying ILS’s cardioprotective effects, transcriptomic analysis was conducted to identify potential targets (Figure 7A-B). By comparing the ISO and ISO+ILS groups, we focused on the aberrant expression of COX6A2, which was significantly downregulated in the ISO group but upregulated in the ISO+ILS group following ILS intervention. Therefore, we hypothesize that the ILS may target COX6A2 and exert cardioprotective effects. Western blot analysis confirmed this hypothesis, showing markedly increased COX6A2 expression in the ISO+ILS group compared to the ISO group (Figure 7C-D).

**Figure 7.**
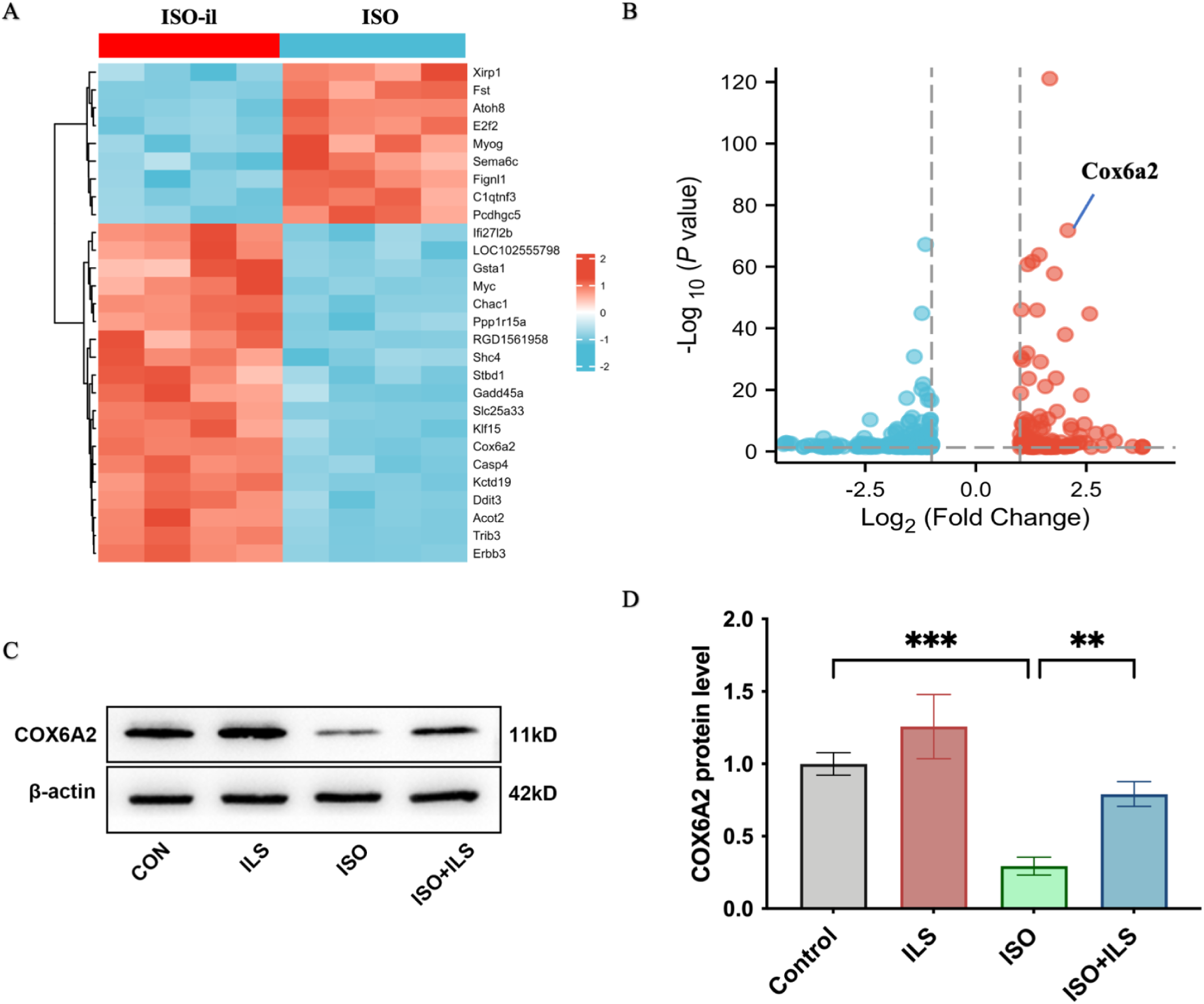
Transcriptomic and biochemical validation of COX6A2 as a target of isoliensinine. (A) Heatmap of differentially expressed genes (DEGs) between ISO and ISO+ILS groups. Rows represent genes (log2 fold change >1.5, p < 0.05); columns represent biological replicates (n = 3/group). COX6A2 (highlighted) was among the most significantly upregulated genes in the ISO+ILS group. (B) Volcano plot of DEGs. COX6A2 (red dot) was identified as a key upregulated gene in the ISO+ILS group. (C-D) Western blot analysis of COX6A2 protein expression. Data presented as mean ± SEM (n ≥ 3), *\*P <* 0.05; *\*\*P <* 0.01; *\*\*\*P <* 0.001; *\*\*\*\*P <* 0.0001.

To confirm COX6A2’s critical role in isoliensinine-mediated protection, siRNA knockdown of COX6A2 was performed (Figure 8A-B). Phalloidin staining revealed no significant change in cell surface area between the si-COX6A2+ISO+ILS group and the si-COX6A2+ISO group, whereas the si-COX6A2+ISO+ILS group exhibited significantly larger cell areas than the si-NC+ISO+ILS group, indicating that COX6A2 knockdown attenuated ILS effects of anti-hypertrophic (Figure 8C-D). Assessment of oxidative stress levels showed that ROS levels in the si-COX6A2+ISO+ILS group remained unchanged compared to the si-COX6A2+ISO group but were significantly higher than those in the si-NC+ISO+ILS group, suggesting that COX6A2 silencing impaired isoliensinine’s antioxidative capacity (Figure 8E-F). Western blot analysis of apoptosis-related proteins demonstrated that the si-COX6A2+ISO+ILS group had upregulated Caspase3 and Bax expression and downregulated Bcl-2 expression compared to the si-NC+ISO+ILS group, indicating that COX6A2 knockdown suppressed isoliensinine’s anti-apoptotic effects (Figure 8G-J). These results collectively demonstrate that isoliensinine protects cardiomyocytes, at least partially, by targeting COX6A2.

**Figure 8.**
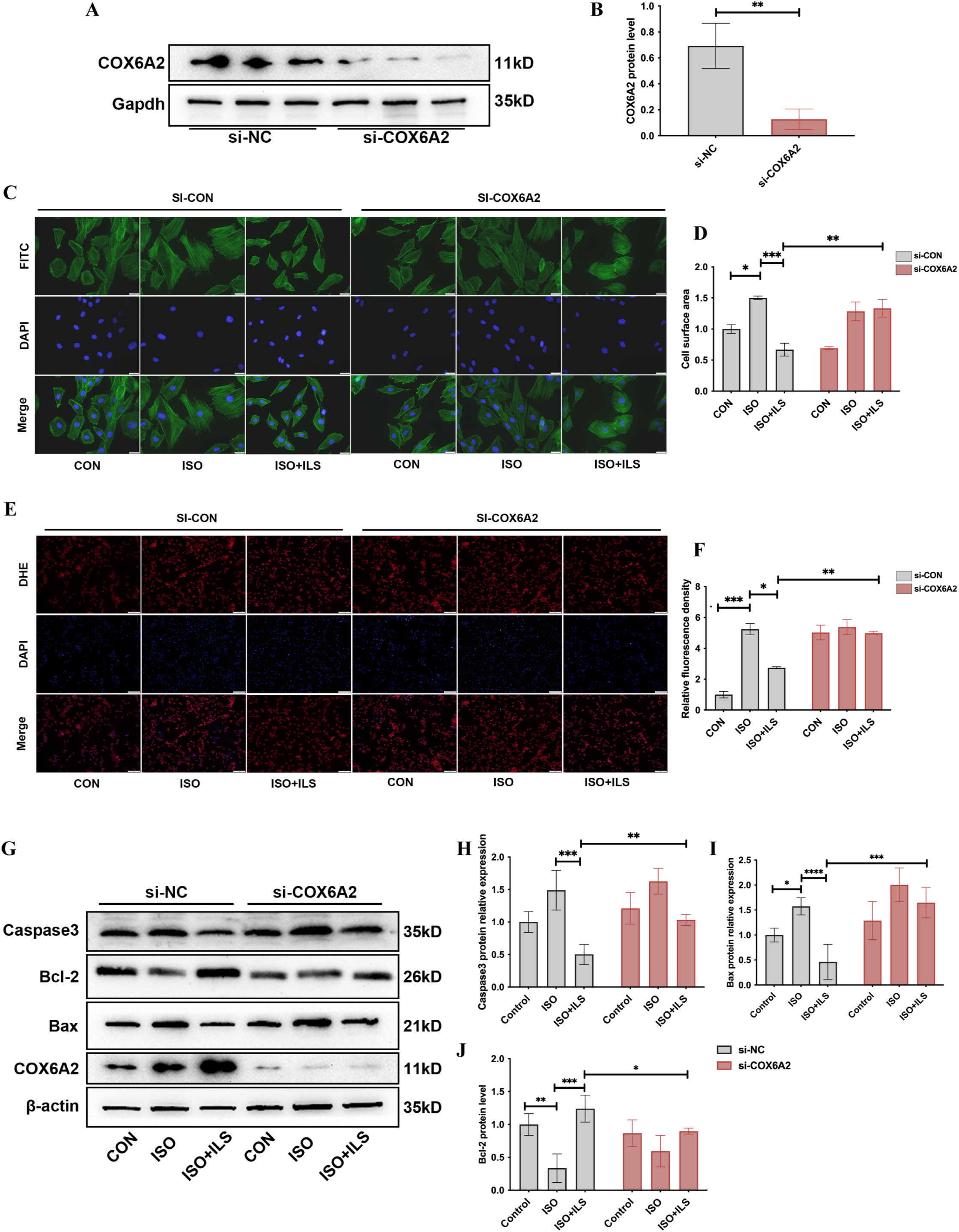
COX6A2 knockdown abolishes isoliensinine-mediated protection against ISO-induced hypertrophy, oxidative stress, and apoptosis. (A-B) Western blot validation of COX6A2 knockdown efficiency. COX6A2 expression was significantly reduced in si-COX6A2-transfected cells compared to si-NC group. (C-D) Phalloidin staining (gree: F-actin; blue: nuclei) and quantitative analysis of cell surface area. Scale bars: 20 μm. (E-F) Fluorescence imaging and quantification of intracellular ROS (DHE probe, red fluorescence). Scale bars: 50 μm. (G-J) Western blot analysis of apoptosis-related proteins: Caspase3, Bax, Bcl-2, and β-actin (loading control). Data presented as mean ± SEM (n ≥ 3), *\*P <* 0.05; *\*\*P <* 0.01; *\*\*\*P <* 0.001; *\*\*\*\*P <* 0.0001.

## 4. Discussion

This study systematically reveals the mechanism by which swimming training ameliorates pathological cardiac hypertrophy by way of metabolic reprogramming (along the isoliensinine-COX6A2 axis). Specifically, swimming training significantly attenuates TAC-induced myocardial hypertrophy, apoptosis, and contractile dysfunction while restoring mitochondrial respiratory function. Integrated metabolomic and transcriptomic analyses demonstrate that swimming training upregulates isoliensinine expression, which targets COX6A2 to suppress oxidative stress, fibrosis, and apoptosis, thereby protecting cardiomyocytes from TAC- or ISO-induced injury.

Extensive studies have confirmed that exercise training, particularly aerobic exercise, exerts significant protective effects against cardiovascular diseases ^20^. Long-term practice indicates that regular aerobic exercise improves cardiovascular function through dual mechanisms of metabolic regulation and structural remodeling ^21,22^. On one hand, chronic aerobic exercise reduces cardiac pressure overload, enhances myocardial contractility, and promotes blood circulation, thereby significantly improving cardiac function ^23,24^. On the other hand, swimming not only increases myocardial oxygen supply but also facilitates mitochondrial biogenesis and functional recovery in the heart, maintaining metabolic homeostasis and meeting energy demands, thereby alleviating pathological cardiac hypertrophy ^25^. Consistent with these findings, our study further supports the notion that swimming serves as an effective intervention to mitigate pathological cardiac hypertrophy. In our results, mice pretreated with swimming training followed by TAC surgery exhibited significantly improved TAC-induced cardiac remodeling and dysfunction, along with attenuated oxidative stress and preserved mitochondrial respiratory function.

Fortunately, our study identifies a potential bridge between aerobic exercise and cardioprotection by revealing a novel molecular mechanism: swimming ameliorates pathological cardiac hypertrophy through the isoliensinine-COX6A2 metabolic signaling axis. Metabolomic profiling revealed that swimming training significantly upregulated endogenous isoliensinine levels in mice. Isoliensinine, a natural alkaloid extracted from the lotus plumule (Nelumbo nucifera), exhibits diverse pharmacological properties, including antioxidant, anti-inflammatory, anti-proliferative, and anti-tumor activities, making it a compound of increasing interest ^26,27^. Previous studies have demonstrated that isoliensinine mitigates inflammatory responses and cellular damage by suppressing inflammatory mediator release and reducing free radical generation, highlighting its broad bioactivity in modulating immune responses and counteracting oxidative stress ^28^. It has also shown therapeutic potential in diseases such as breast cancer ^29^ and hypertension ^30^ through multi-pathway regulation.

However, the role of isoliensinine in cardiovascular diseases, particularly pathological cardiac hypertrophy, remains underexplored, and its cardioprotective mechanisms are poorly understood. Our study is the first to reveal the critical role of isoliensinine in regulating cardiac hypertrophy. We demonstrated that exogenous isoliensinine administration significantly attenuated TAC or ISO induced myocardial hypertrophy, interstitial fibrosis, apoptosis, and oxidative stress. This novel discovery expands our understanding of isoliensinine’s biological functions and highlights its therapeutic potential for cardiovascular disorders.

Further mechanistic investigations suggest that ILS’s cardioprotective effects may stem from its synergy with mitochondrial function. As the central hub of cellular energy metabolism, mitochondria not only supply ATP essential for cardiomyocytes but also regulate redox balance, calcium homeostasis, and apoptosis ^13,31^. Previous studies have highlighted the pivotal role of mitochondrial dysfunction in the initiation and progression of cardiac hypertrophy ^12,32^. Our transcriptomic analysis is the first to directly link COX6A2 to ILS-mediated attenuation of cardiac remodeling. COX6A2, a critical subunit of mitochondrial electron transport chain complex IV, regulates oxidative phosphorylation and ATP production. Studies indicate that COX6A2 deficiency leads to increased reactive oxygen species (ROS) generation ^33^. In pathological cardiac hypertrophy, the heightened metabolic demands of cardiomyocytes coincide with mitochondrial dysfunction, including disrupted biogenesis, elevated oxidative stress, and structural/functional decompensation ^34,35^. Notably, ISO treatment significantly downregulated COX6A2 expression, aligning with observed mitochondrial ultrastructural damage and oxidative stress during pathological hypertrophy. ILS counteracted these effects by upregulating COX6A2, thereby mitigating oxidative stress and cardiomyocyte injury. This conclusion was further validated by siRNA-mediated COX6A2 knockdown, which abolished ILS’s cytoprotective effects. Through this regulatory mechanism, ILS effectively resisted pressure overload (TAC)- or ISO-induced cardiomyocyte hypertrophy and improved overall cardiac function. These findings highlight COX6A2 as a promising therapeutic target for future research on heart failure.

In summary, our study is the first to demonstrate that swimming training attenuates ventricular remodeling and dysfunction by modulating the Isoliensinine-COX6A2 signaling axis, enhancing mitochondrial function, and promoting energy metabolic homeostasis. These findings not only deepen our understanding of the molecular mechanisms underlying aerobic exercise (particularly swimming) in alleviating cardiac hypertrophy but also identify novel potential therapeutic targets for cardiovascular diseases. The regulatory pathway involving ILS and COX6A2 holds significant promise for the prevention and treatment of heart failure. However, the upstream regulatory networks of ILS (e.g., how swimming activates its biosynthesis) remain unelucidated. Additionally, whether COX6A2 interacts with other cardioprotective pathways warrants further investigation. Future studies will focus on elucidating the mechanistic roles of ILS and COX6A2 in pathological cardiac hypertrophy and validating their broad applicability across diverse pathological conditions, providing new perspectives and therapeutic targets for cardiac protection.

## Sources of Funding

This research is supported by the National Natural Science Foundation of China [Grant Nos. 81860086, 82400462, 82300434], the Natural Science Foundation of Jiangxi Province [Grant No. 20242BAB21039], and the “Yangfan Project” for Talent Cultivation of the First Affiliated Hospital of Nanchang University.

## Disclosures

None.

## Abbreviations

ISO: isoproterenol
TAC: transverse aortic constriction
COX6A2: Cytochrome C Oxidase Subunit 6A2
ILS: isoliensinine

## Notes

**Conflict of Interest Statement:** The authors have no conflicts of interest to declare.

### Competing Interest Statement

The authors have declared no competing interest.

